# *N*-acetylglucosamine supplementation fails to bypass the critical acetylation of glucosamine-6-phosphate required for *Toxoplasma gondii* replication and invasion

**DOI:** 10.1101/2024.01.18.576165

**Authors:** María Pía Alberione, Víctor González-Ruiz, Serge Rudaz, Dominique Soldati-Favre, Luis Izquierdo, Joachim Kloehn

## Abstract

The cell surface of *Toxoplasma gondii* is rich in glycoconjugates which hold diverse and vital functions in the lytic cycle of this obligate intracellular parasite. Additionally, the cyst wall of bradyzoites, that shields the persistent form responsible for chronic infection from the immune system, is heavily glycosylated. Formation of glycoconjugates relies on activated sugar nucleotides, such as uridine diphosphate *N*-acetylglucosamine (UDP-GlcNAc). The Glucosamine-phosphate-*N*-acetyltransferase (GNA1) generates *N*- acetylglucosamine-6-phosphate critical to produce UDP-GlcNAc. Here, we demonstrate that downregulation of *T. gondii* GNA1 results in a severe reduction of UDP-GlcNAc and a concomitant drop in glycosylphosphatidylinositol (GPI), leading to impairment of the parasite’s ability to invade and replicate in the host cell. Surprisingly, attempts to rescue this defect through exogenous GlcNAc supplementation fail to completely restore these essential functions. In depth metabolomic analyses elucidate diverse causes underlying the failed rescue: utilization of GlcNAc is inefficient under glucose-replete conditions and fails to restore UDP-GlcNAc levels in GNA1-depleted parasites. In contrast, GlcNAc- supplementation under glucose-deplete conditions fully restores UDP-GlcNAc levels but fails to rescue the defects associated with GNA1 depletion. Our results underscore the essentiality of GlcN6P acetylation in governing *T. gondii* replication and invasion and highlight the potential of the evolutionary divergent GNA1 in Apicomplexa as a target for the development of much-needed new therapeutic strategies.

## Introduction

The phylum of Apicomplexa groups a vast number of obligate intracellular parasites, some of which pose a considerable threat to human health. The most ubiquitous apicomplexan, *Toxoplasma gondii*, causes disease in immunocompromised individuals [1, 2], as well as abortions, stillbirths, fetal death, retinal lesions or long-term disabling sequelae in congenitally infected children [3, 4]. At present, there is no vaccine that prevents toxoplasmosis, and the available treatments are associated with a range of shortcomings including high cost, toxicity and rising resistance [5]. In the accidental human host, *T. gondii* manifests in two distinct stages: the fast-replicating tachyzoite, responsible for acute disease and the slow replicating bradyzoite, which persists encysted within muscle cells and neurons throughout the lifetime of its host [6]. These persistent parasites constitute a reservoir, that can reactivate causing life-threatening acute toxoplasmosis when the infected individual becomes immunocompromised. The inability to eradicate the parasite’s latent form, combined with the emergence of parasites that are resistant to existing drugs against acute toxoplasmosis, underscores the pressing need for novel therapeutic strategies [5].

The endomembrane system of *T. gondii* is rich in glycoconjugates which play fundamental roles in infectivity, survival, and virulence [7]. Several glycan structures have been characterized in *T. gondii* including *N*-glycans [8], *O*-glycans [9-13], *C*- mannose [9, 14], GPI-anchors [15, 16], and others [7]. These glycans serve various critical functions from invasion to O2 sensing and nutrient storage, hence contributing to the overall virulence of the parasite [7]. Additionally, glycans are critical components of the bradyzoite cyst wall and the disruption of their formation impairs the parasite’s ability to persist [17, 18].

The *de novo* synthesis of glycans relies on activated sugar nucleotides. Uridine diphosphate *N*-acetylglucosamine (UDP-GlcNAc) serves as a donor by GlcNAc-dependent glycosyltransferases for the synthesis of *N-*glycans, glycosylphosphatidylinositol (GPI)-anchors, glycoinositolphospholipids (GIPLs), and for the glycosylation of other protein acceptors. Given the critical roles of these structures for infectivity of tachyzoites [7, 8, 16, 19], and bradyzoite survival and replication [17, 18], the biosynthesis route of UDP-GlcNAc is a plausible target for intervention against acute toxoplasmosis and for eradication the chronic infection. GNA1, the enzyme catalysing the acetylation of glucosamine-6-phosphate (GlcN6P) is considered a promising drug target in Apicomplexa. This is attributed to its independent evolutionary origin, unique sequence features [20], and established essentiality for the intraerythrocytic development of *Plasmodium falciparum* [21].

In *T. gondii*, a genome-wide CRISPR fitness screen underscored the significance of UDP-GlcNAc biosynthesis for the parasite, classifying several genes encoding for enzymes involved in the amino sugar synthesis pathway as fitness-conferring [22, 23]. Unexpectedly, however, this study predicted GNA1 to be dispensable for *T. gondii*, even though the upstream and downstream enzymes were highly fitness-conferring [23].

Here we demonstrate the essential nature of GNA1 in *T. gondii*, revealing that its downregulation leads to a reduction in GPI anchors that impairs invasion and replication within its host cell. Intriguingly, defects in GNA1 cannot be overcome by GlcNAc salvage. Targeted metabolomic analyses revealed that GlcNAc salvage is inefficient in glucose-replete conditions. In contrast, GlcNAc is effectively salvaged in glucose-deplete conditions, but fails to rectify the defects associated with the disruption of the pathway. These findings highlight the potential of GNA1 as a drug targe for combatting toxoplasmosis.

## Results

### *T. gondii* GNA1 is essential, contradicting the prediction from a genome wide fitness screen

*T. gondii* expresses several glycan structures akin to other eukaryotic cells, with few noteworthy characteristics (Fig 1A). The synthesis of GPI-anchors, *N*-glycans and *O*- glycans requires as donor substrate UDP-GlcNAc, or UDP-GalNAc. The latter can be synthesised from UDP-GlcNAc via GalE [7], a UDP-Glc/UDP-Gal epimerase. In the amino sugar biosynthesis pathway, the glucosamine 6-phosphate *N*-acetyltransferase,

**Fig 1.**
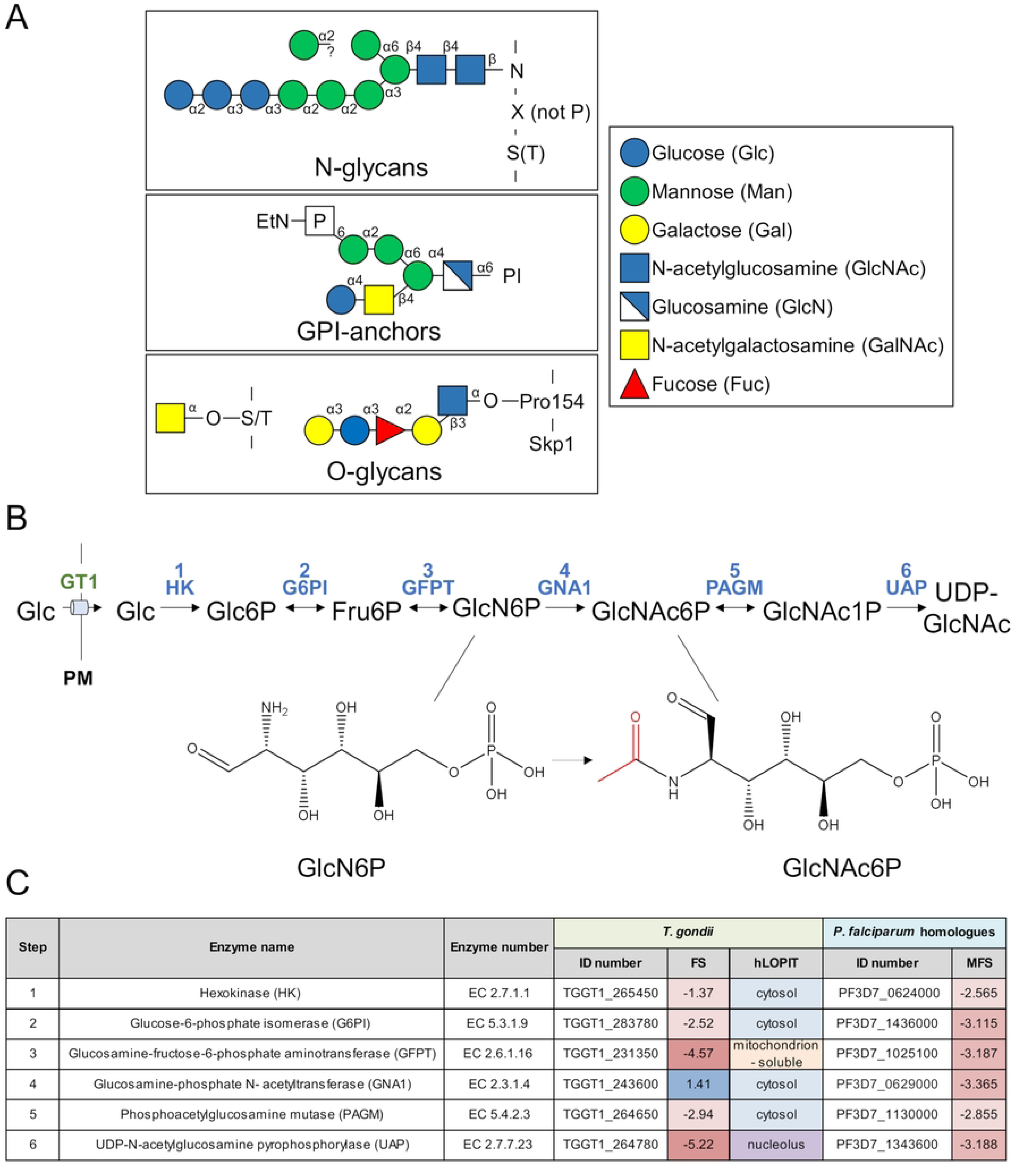
Glycans and UDP-GlcNAc synthesis in *T. gondii*. A) Proteins of the secretory pathway of *Toxoplasma gondii* are commonly modified as *N*-glycans, *O*-glycans or GPI- anchored. The typical glycan structures in *T. gondii* are shown, as well as the modification of Skp1, which harbours a specific *O*-glycosylation [65]. **B)** The activated sugar nucleotide UDP-GlcNAc is synthesised from glucose in six conserved enzymatic reactions, with the glucosamine-phosphate *N*-acetyltransferase (GNA1) converting glucosamine-6-phosphate (GlcN6P) to *N*-acetylglucosamine-6-phosphate (GlcN6P) **C)** Overview of *T. gondii* enzymes in UDP-GlcNAc synthesis, listing their names, accession number (ID) [25], fitness score (FS) [23] and putative localisation (hLOPIT) [66], as well as the accession number [67] and mutagenesis fitness score (MFS) [68] in the related *Plasmodium falciparum* parasite. Abbreviations: PI: phosphatidylinositol; EtN: ethanolamine; GT1: glucose transporter 1; PM: plasma membrane; Fru, fructose; PI, phosphatidylinositol; GPI, glycosylphosphatidyl inositol; UDPGlcNAc, uridine diphosphate *N*-acetylglucosamine. Other abbreviations, see panel A and C.

GNA1, catalyses the acetylation of GlcN6P (Fig 1B). Although *T. gondii* has been shown to critically rely on several glycan structures [7-9], *Tg*GNA1 (TGGT1_243600) was assigned a positive fitness score (+1.41) in a genome-wide fitness screen [23], indicating its potential dispensability (Fig 1C). This is unexpected considering the crucial role of GNA1 in other organisms [24], including *P. falciparum* [21]. Additionally, enzymes acting either downstream or upstream of GNA1 were assigned negative fitness scores in *T. gondii*, suggesting their essentiality [23].

Examination of the nanopore sequencing data on ToxoDB [25, 26] revealed two major GNA1 transcripts, with the shorter, more prevalent transcript only covering a portion of the predicted protein coding sequence. In addition to the predicted GNA1 protein coding sequence, which encodes a putative protein of 55.5 kDa, four additional in-frame open reading frames were identified. These could give rise to GNA1 proteins of various reduced sizes (45.5, 21.0, 17.2 and 16.7 kDa) [25] (S1 Fig). Critically, the acetyltransferase domain is located near the C-terminus and is present in all five putative isoforms. Consequently, all proteins, including the shortest version, could potentially be catalytically active. These shorter GNA1 isoforms may explain the unexpected positive fitness score for GNA1 [23]. Concordantly, out of the ten single guide RNAs (sgRNAs) used to target GNA1 in the genome wide fitness screen [25, 26], four bind near the extended N-terminus, only present in the longest isoform of GNA1 (guides sgTGGT1_243600_5, _6, _9 and _10; average fitness score +1.02), while three bind close to the C-terminus of GNA1, disrupting all potential isoforms, (guides sgTGGT1_243600_2, _3, and _4; average fitness score: ™7.50,) (S1 Fig). Notably, the published phenotype score is calculated by averaging the value of the top five scoring guides to mitigate the impact of stochastic losses, resulting in the positive fitness score for GNA1 [23]. Given the existence of diverse isoforms, the assigned fitness score likely does not reflect the importance that GNA1 may play for *T. gondii*, prompting further investigation of the enzyme.

To examine the localization and function of GNA1 in *T. gondii* (TGGT1_243600), the endogenous locus was edited using CRISPR/Cas9 [23, 27]. Simultaneously a Ty epitope tag and a mini auxin inducible degron (mAID) domain were fused to the C-terminus of GNA1 in parasites stably expressing the auxin receptor transport inhibitor response 1 (TIR1) from *Oryza sativa* (RH-TIR1) [28, 29]. The hypoxanthine-xanthine-guanine phosphoribosyl transferase (*hxgprt*) resistance cassette was inserted, for selection of positive transfectants (S1 Fig) [30]. The mAID domain enables rapid and efficient downregulation of the protein of interest via proteasomal degradation upon addition of auxin (indole 3-acetic acid, IAA) [29]. Successful integration of the construct at the *gna1* locus in a clonal population was confirmed by genomic PCR (S1 Fig), using primers listed in S2 Table. GNA1-mAID-Ty exhibited a dotty cytosolic localization by immunofluorescence assay (IFA) (Fig 2A). Subsequently, downregulation of GNA1 was assessed by Western blot (2, 4- and 18-hours IAA treatment, Fig 2B) and by IFA (18-hours IAA treatment, Fig 2C), confirming an efficient and complete depletion of GNA1 after 2-4 hours of IAA treatment. Interestingly, the western blot revealed several bands between approximately 30-80 kDa for GNA1-mAID-Ty (Fig 2B), all of which were efficiently downregulated upon addition of IAA. Up to five GNA1 isoforms may exist, with molecular weights ranging from 29.7 – 68.5 kDa, including the tag. These results indicate that full length GNA1 as well as shorter isoforms (S1 Fig) are synthesised, although partial degradation cannot be excluded.

**Fig 2.**
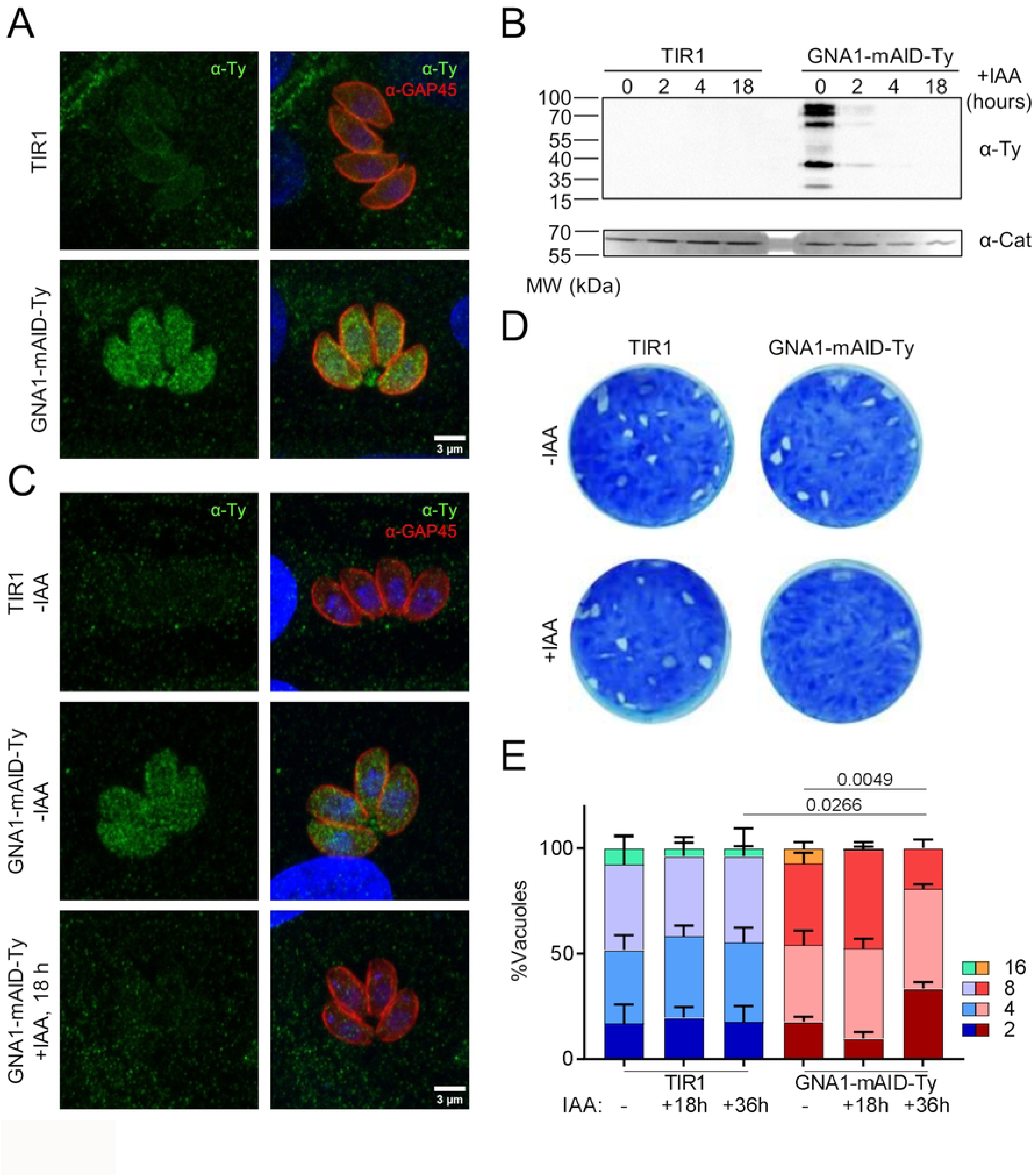
GNA1 is essential for *T. gondii*. A) Immunofluorescence assay (IFA) showing the pellicle marker GAP45 and Ty signal in GNA1-mAID-Ty parasite line and its parental line (TIR1). **B)** Western blot revealing the signal of Ty-tagged GNA1 and TIR1 at different time points of auxin (IAA) treatment. **C)** IFA showing Ty signal in GNA1-mAID-Ty parasites in the absence of IAA and 18 hours after IAA treatment. **D)** Lysis plaques formed over one week of TIR1 and GNA1-mAID-Ty parasite cultivation in the presence or absence of IAA. **E)** Growth assay of TIR1 and GNA1-mAID-Ty parasites showing the number of parasites per vacuole after 24 hours of growth and varying durations of IAA treatment. A-D show representative data of 3 independent experiments. E shows representative data from one of three independent biological replicates averaging technical triplicates. p-values are given comparing the average number of parasites by two-sided student’s t-test, between the indicated conditions. Abbreviations: MW, molecular weight; CAT, catalase.

The significance of GNA1 for the parasite lytic cycle was assessed by plaque assay. Downregulation of GNA1 prevented the formation of plaques of lysis (Fig 2D), indicating that GNA1 is needed for one or several steps of the lytic cycle. The intracellular growth assay revealed a significant impact of GNA1 depletion on the replication rate, with the average number of parasites during 24 hours of growth decreasing from 6.5 in the controls to 4.2 following 36 hours of IAA treatment (Fig 2E).

These results underscore the essential role of GNA1 in intracellular growth and overall lytic cycle of *T. gondii,* consistent with the importance of glycoconjugates. The data suggest that two major transcripts are generated for GNA1, yielding up to five protein isoforms that can be potentially catalytically active. The extended N-terminus in the longer isoform may be dispensable [23]. If it holds a regulatory function in other life cycle stages remains unknown. Crucially, depletion of the GNA1 acetyltransferase domain is detrimental to *T. gondii*.

### GNA1 is active and critical for UDP-GlcNAc synthesis in *T. gondii*

To examine if the UDP-GlcNAc biosynthesis pathway is active in intra-and extracellular *T. gondii*, TIR1 parasites were cultured intracellularly for 24 hours in medium containing 10 mM uniformly ^13^C-labelled glucose (U-^13^C6-Glc) or extracted and purified extracellular parasites were incubated for 3 hours in medium containing heavy Glc. Post-incubation, the metabolism was quenched, parasites were harvested, and metabolites extracted. Gas chromatography-mass spectrometry (GC-MS) following derivatization of the sugars was employed to assess the extent of label incorporation into *N*-acetylglucosamine-6-phosphate (GlcNAc6P) (S3 Fig). Synthesis of GlcNAc6P from labelled Glc was observed in both parasite stages, reaching 88.8% and 25.5% labelling in intra-and extracellular parasites, respectively (Fig 3A). These findings unequivocally demonstrate the active UDP-GlcNAc biosynthesis pathway from Glc in both intra-and extracellular *T. gondii*.

**Fig 3.**
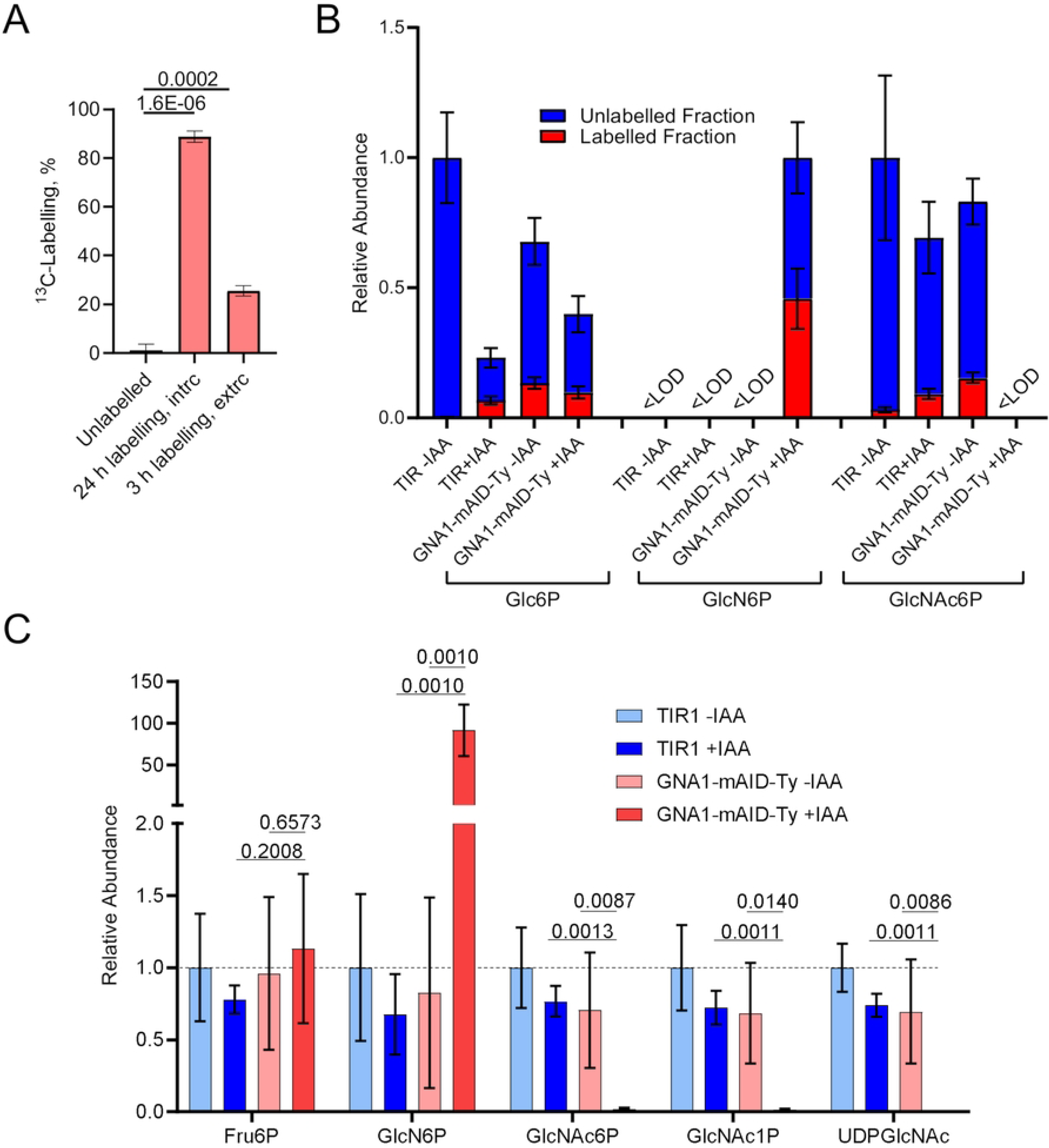
UDP-GlcNAc synthesis is disrupted in *T. gondii* that lack GNA1. **A)** Percent ^13^C-labelling in *T. gondii* (TIR1) derived *N*-acetylglucosamine-6-phosphate (GlcNAc6P) in unlabelled parasites (natural abundance) or after incubation of intracellular or extracellular parasites in medium containing U-^13^C6-glucose for 24 or 3 hours, respectively. **B)** Relative abundance and fractional ^13^C-labelling in TIR1 and GNA1-mAID-Ty parasite metabolite extracts, following incubation of extracellular parasites in medium containing U-^13^C6-glucose in the absence of auxin (−IAA) or following 18 hours pre-treatment (+IAA). TIR1 −IAA parasites were incubated in medium with natural abundance glucose as an unlabelled control. Note that metabolites for which the abundance of labelled and unlabelled ions was too low to obtain reliable labelling data were deemed below limit of detection (<LOD, sum of ion intensity <1000 arbitrary units). **C)** Relative metabolite levels in TIR1 and GNA1-mAID-Ty, following no treatment (−IAA) or treatment with IAA for 18 hours during intracellular growth (+IAA). Data plotted show the average and standard deviation of 3 (A, B) or 4 (C) independent biological replicates. p-values from two-sided student t-tests are given in A and C, comparing the indicated conditions. Abbreviations: Glc6P, glucose-6-phosphate; GlcN6P, glucosamine-6-phosphate; GlcNAc6P, *N*-acetylglucosamine-6-phosphate; Fru6P, fructose-6-phosphate; GlcNAc1P, *N*-acetylglucosamine-1-phosphate; UDPGlcNAc, uridine diphosphate *N*-acetylglucosamine.

To assess the critical participation of GNA1 in the pathway, TIR1 and GNA1-mAID-Ty parasites were untreated or pre-treated with IAA for 18 hours and extracellular parasites were incubated in medium containing 10 mM U-^13^C6-Glc. TIR1 −IAA parasites that served as control, were incubated with regular medium containing unlabelled (natural abundance) Glc. After 5 hours of incubation, parasites were harvested, metabolites extracted, derivatized and analysed by GC-MS in a targeted manner (S3 Fig). Under all tested conditions, Glc6P was detected with incorporation of heavy carbons under ^13^C-labelling conditions (Fig 3B). Intriguingly, glucosamine-6-phosphate (GlcN6P), the substrate of GNA1, was exclusively detected in parasites depleted in GNA1, with incorporation of considerable labelling (45.8%). In contrast, GlcNAc6P, the product of GNA1, was detected in all conditions, except in parasites depleted in GNA1 (Fig 3B). To gain a comprehensive understanding of the impact of GNA1 depletion on *T. gondii* metabolism, we performed metabolite profiling by GC-MS after 36 hours of downregulation (S4 Fig). At this relatively late time point, pleiotropic effects were observed with 28 out of 64 metabolites significantly altered (>2-fold) in their abundance compared to TIR1 −IAA. The majority (26 metabolites) exhibited reduced abundance including amino acids, TCA cycle intermediates, sugars, fatty acids, and others. Amongst the most dramatically reduced metabolites were GlcNAc and GlcNAc6P, consistent with the function of GNA1. Conversely the two significantly increased metabolites were GlcN and myo-inositol. While GlcN accumulation directly correlates with the absence of GNA1 (following enzymatic dephosphorylation or loss of the phosphate group during sample preparation from GlcN6P), accumulation of myo-inositol could be part of a general stress response, as previously observed [31], or a consequence of impaired GPI-anchor synthesis.

Given that crucial intermediates such as GlcNAc1P and the product UDP-GlcNAc cannot be detected by GC-MS, we turned to liquid chromatography mass spectrometry (LC-MS/MS) for more sensitive detection of all relevant pathway intermediates. Intracellular parasites were treated with IAA or not for 18 hours, before quenching of the metabolism, parasite harvest, metabolite extraction and analysis. While fructose-6-phoshate (Fru6P) levels remained unaffected by GNA1 downregulation (GNA1-mAID-Ty +IAA), GlcN6P accumulated dramatically to 91.8-fold higher levels compared to the controls (Fig 3C). In sharp contrast, as observed by GC-MS, GlcNAc6P was markedly reduced (50.9-fold). Similarly, the subsequent metabolites, GlcNAc1P and the product UDP-GlcNAc exhibited reductions of 83.3- and 803.2-fold, respectively (Fig 3C).

Together, these findings reveal that UDP-GlcNAc synthesis is active both in intra-and extracellular *T. gondii* tachyzoites, Moreover, GNA1 plays a critical role in this pathway as its disruption results in a significant reduction in the activated sugar nucleotide UDP-GlcNAc and an accumulation of its substrate GlcN6P.

### GlcNAc supplementation fails to rescue GNA1 deficiency

Efficient bypass of defects in UDP-GlcNAc synthesis is well documented in several organisms, including *P. falciparum* through GlcNAc supplementation [21, 32, 33]. GlcNAc can be taken up and phosphorylated by hexokinase, generating GlcNAc6P, effectively circumventing the initial steps of the pathway. To test if exogenous GlcN or GlcNAc supplementation can bypass the function of GNA1 in *T. gondii*, we performed plaque assays with GNA1-mAID-Ty parasites in presence or absence of IAA while supplementing different concentrations of GlcN or GlcNAc. Remarkably, none of the supplementations could rescue the lytic cycle defect observed in parasites depleted in GNA1 (Fig 4A).

**Fig 4.**
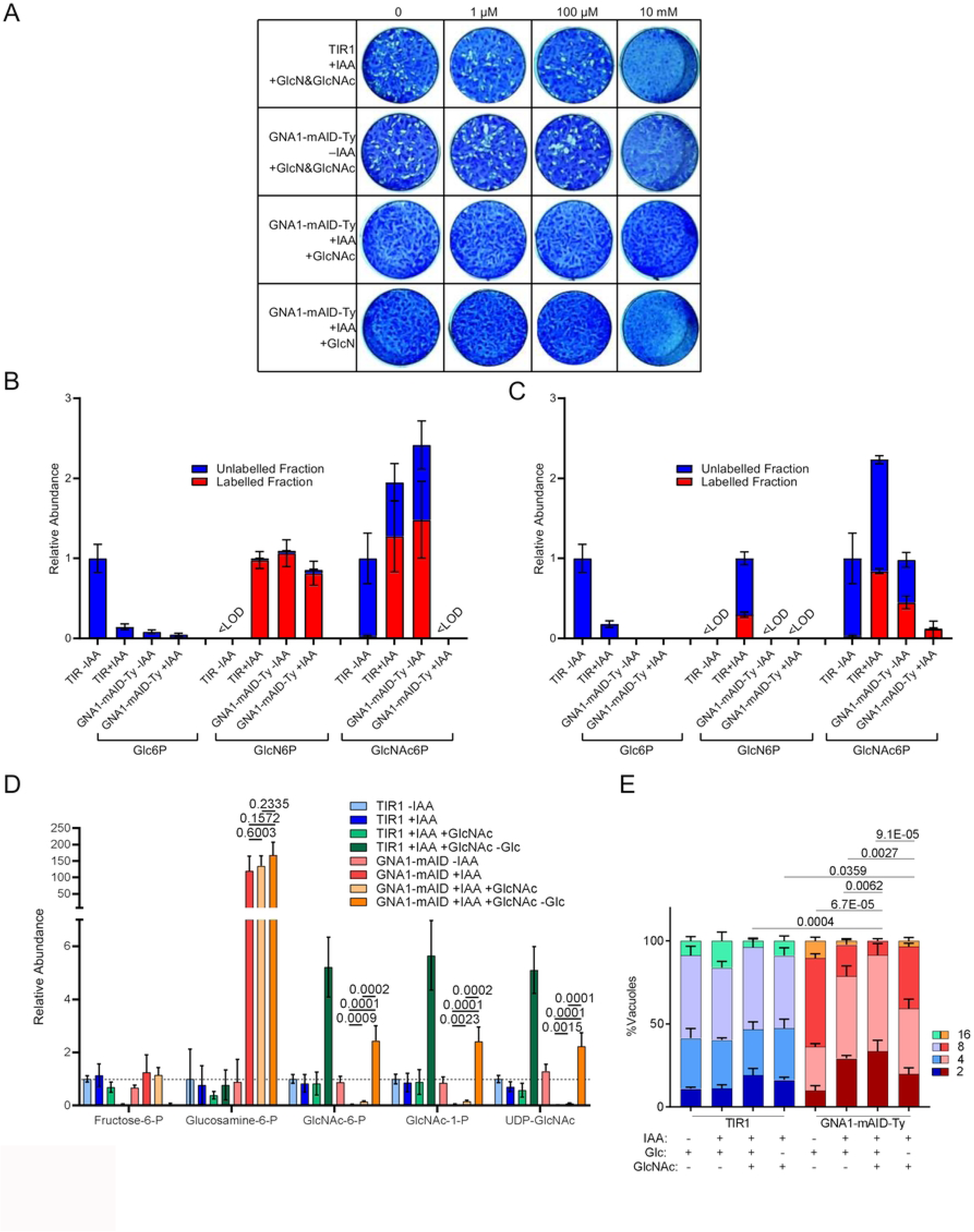
Disruption of *Tg*GNA1 cannot be rescued by GlcNAc supplementation. **A)** Lysis plaques formed by TIR1 and GNA1-mAID-Ty parasites over one week of growth when treated with auxin (+IAA) or not (−IAA) and supplemented with varying concentrations of glucosamine (GlcN) and/or *N*-acetylglucosamine (GlcNAc) as indicated. **B-C)** Relative metabolite abundance and fractional ^13^C-labelling in TIR1 and GNA1-mAID-Ty parasite extracts, incubated for 5 hours extracellularly in medium without glucose and containing U-^13^C6-glucosamine (**B**) or U-^13^C6-*N*-acetylglucosamine (**C**) in the absence of IAA or following IAA pre-treatment (+IAA, 18 h). TIR1 −IAA parasites were incubated in medium with natural abundance glucose as an unlabelled control. Note that metabolites for which the abundance of labelled and unlabelled ions was too low to obtain reliable labelling data were deemed below limit of detection (<LOD, sum of ion intensity <1000 arbitrary units). **D)** Relative metabolite levels in TIR1 and GNA1-mAID- Ty, following no treatment (−IAA) or treatment with IAA (+IAA, 18 h) during intracellular growth in the presence or absence of glucose and supplemented with GlcNAc as indicated for the same duration. **E)** Intracellular growth assay showing the number of parasites per vacuole after 24 hours of growth, treated for 48 hours with IAA, Glc or GlcNAc as indicated. A) shows representative images of 3 independent experiments. Data plotted in B-E show the average and standard deviation of 3 (B, C, E) and 4 (D) independent biological replicates, respectively. p-values from two-sided student t-tests are given in D and E, comparing the indicated conditions. p-values in E compare the average number of parasites per vacuole. Abbreviations: Glc, glucose. Other abbreviations, see Fig 3.

We hypothesised that the inability of GlcN or GlcNAc supplementation to rescue the lytic cycle defect in GNA1-depleted parasites could be attributed to various reasons I) insufficient uptake of GlcNAc by the host cells and/or *T. gondii*, II) incapacity of *T. gondii* hexokinase to phosphorylate GlcNAc or III) inefficient entry of phosphorylated, salvaged GlcNAc into the UDP-GlcNAc synthesis pathway. To explore these possibilities, we incubated purified extracellular parasites in medium without Glc supplemented with ^13^C6- GlcN or ^13^C6-GlcNAc for 5 hours, before harvesting parasites and extracting metabolites. GNA1-mAID-Ty and TIR1 parasites were pretreated with IAA for 18 hours to deplete GNA1 levels in the GNA1-mAID-Ty strain. TIR1 −IAA were incubated in regular medium with unlabelled (natural abundance) Glc. Remarkably, both amino sugars were efficiently salvaged and utilised. ^13^C6-GlcN was salvaged and phosphorylated under all conditions, but as expected, the downregulation of GNA1 prevented the formation of GlcNAc6P from GlcN6P (Fig 4B). Similarly, exogenous ^13^C6-GlcNAc was efficiently used to generate labelled GlcNAc6P in the control strains (Fig 4C). Parasites deficient in GNA1 could also salvage exogenous ^13^C6-GlcNAc and utilised it to generate GlcNAc6P, albeit at significantly lower levels but fully ^13^C-labelled, consistent with the inability of Glc to contribute to GlcNAc6P formation. These results reveal that *T. gondii* can salvage and utilise GlcNAc, potentially bypassing the need for GNA1.

It is noteworthy that this experiment was conducted with extracellular parasites which were pre-depleted in GNA1 over 18 hours under regular growth conditions, without GlcNAc supplementation. Thus, parasites were expected to be impaired in their fitness and this measurement may not reflect what occurs during intracellular development and during continuous supplementation. To address this, we cultured intracellular *T. gondii*, TIR1 and GNA1-mAID-Ty parasites under standard conditions for 24 hours before changing the culture medium to one of the following 4 conditions for 18 hours prior to parasite harvest: regular medium −IAA; regular medium +IAA; regular medium +IAA supplemented with 10 mM GlcNAc; and medium without Glc +IAA supplemented with additional glutamine and 10 mM GlcNAc. Parasites were harvested while intracellular, with the medium being removed and parasite metabolism quenched prior to the harvest of parasites, to exclude any metabolite uptake by extracellular parasites. Metabolites were extracted and analysed by LC-MS/MS, as above (Fig 4D). Remarkably, GlcNAc supplementation had only a marginal impact in the presence of Glc. Although significantly higher than in non-supplemented parasites devoid of GNA1, GlcNAc failed to fully restore UDP-GlcNAc levels, with levels being 13.3-fold lower than in control (TIR1-IAA) parasites (see detail in S5 Fig). In the absence of Glc, however, GlcNAc was efficiently salvaged and markedly increased UDP-GlcNAc levels, both in TIR1 parasites as well as in parasites deficient of GNA1. Crucially, UDP-GlcNAc levels in GNA1-devoid parasites supplemented with GlcNAc in the absence of Glc were 2.2-fold higher than in control (TIR1-IAA) parasites, suggesting a full rescue of the pathway. Unexpectedly, GlcN6P continued to accumulate to levels >100-fold higher in parasites devoid of GNA1, regardless of the presence or absence of Glc in the medium. Since Fru6P levels were markedly down in the absence of Glc, we speculate that the detected GlcN6P is not derived from gluconeogenesis but rather from the deacetylation of GlcNAc, either through deacetylases of the host cell or by the parasite. The generated GlcN6P failed to be converted further in the absence of GNA1. In summary, this detailed analysis of the pathway under varying conditions reveals efficient GlcNAc salvage but only in the absence of Glc. Under this condition, the function of GNA1 can be bypassed, fully restoring UDP-GlcNAc levels. Whether a potential competition between Glc and GlcNAc happens at the level of uptake by the host or the parasite, or at the level of phosphorylation in the parasite remains unclear.

While GlcNAc supplementation failed to restore the lytic cycle defect in GNA1-depleted parasites in the presence of Glc, we investigated whether the defects in the parasite’s intracellular growth could be rescued by exogenous GlcNAc, both in the presence or absence of Glc. Consistent with the inefficient utilization of GlcNAc in the presence of Glc, GlcNAc supplementation failed to rescue the growth defect in regular medium (Fig 4E). GlcNAc supplementation in the absence of Glc, however, facilitated a significant but modest and incomplete rescue of the intracellular growth rate, following 48 hours of treatment (Fig 4E). Lastly, we explored whether certain ratios of Glc and GlcNAc could potentially rescue parasites depleted in GNA1, by supporting central carbon metabolism (Glc), but facilitating GNA1 bypass (GlcNAc) and potentially alleviating excessive GlcN6P accumulation. Plaque assays were performed with parasites in varying Glc and GlcNAc ratios, however none of the conditions were able to rescue the lytic cycle defect associated with GNA1 downregulation (S6 Fig).

### GNA1 is needed for GPI-anchor synthesis critical for host cell invasion by *T. gondii*

The observed impairment of UDP-GlcNAc synthesis in parasites depleted in GNA1 is expected to impair the synthesis of glycans, including GPI-anchors, thereby likely affecting the parasite’s ability to invade host cells. GPI-anchored proteins are critical for parasite invasion, contributing to the expression of a series of surface antigens, which play a vital role during host cell attachment [16, 34, 35]. Additionally, glycosylated proteins have been described to traffic to the apical secretory organelles and playing an essential role in their biogenesis and function [19, 36]. To assess if GNA1 is required for the appropriate localization and formation of GPI-anchored proteins, we performed IFAs, evaluating the expression and distribution of the surface antigen 1 (SAG1) [37]. Downregulation of GNA1 resulted in a drop in SAG1 signal intensity and an aberrant distribution, with the signal commonly accumulating in the residual body (Fig 5A). This abnormal staining was observed in 68.3 and 73.3% of vacuoles following downregulation of GNA1 for 18 or 36 hours, respectively (Fig 5B). To investigate if this is a relatively specific defect or if cells devoid of GNA1 exhibit various morphological abnormalities, we assessed the morphology of the apicoplast and mitochondrion, upon GNA1 downregulation for the same duration (S7 Fig). The organelles were studied by IFA using α-CPN60 (chaperonin 60) and α-5F4 (F1 ATPase beta subunit), two specific markers found in the apicoplast and at the mitochondrion, respectively. Both organelles appeared morphologically intact and normal after 18 hours and 36 hours of IAA-treatment (S7 Fig). Next, to confirm if the abnormal SAG1 signal, observed in GNA1-deficient cells, can indeed be attributed to a defect in the synthesis of GPI-anchors, we quantified the relative amount of GPI-anchors following downregulation of GNA1 for 36 hours. To this end, parasite lipids were extracted and subjected to methanolysis to hydrolyse monosaccharides off glycan structures found in the organic phase after metabolite extraction. The derivatised sugar and fatty acid methyl esters were analysed by GC-MS and the signal intensity for mannose residues was quantified relative to the signal intensity for palmitic acid [38]. Notably, the ratio mannose signal intensity/palmitic acid signal intensity decreased 8.5-fold after 36 hours of IAA treatment, consistent with a marked drop in GIPLs and GPI-anchor formation (Fig 5C). As expected, this caused a severe defect in host cell invasion, with only 26.5% of parasites depleted in GNA1 over 18 hours invading successfully, compared to more than 70% of parasites in all controls (Fig 5D).

**Fig 5.**
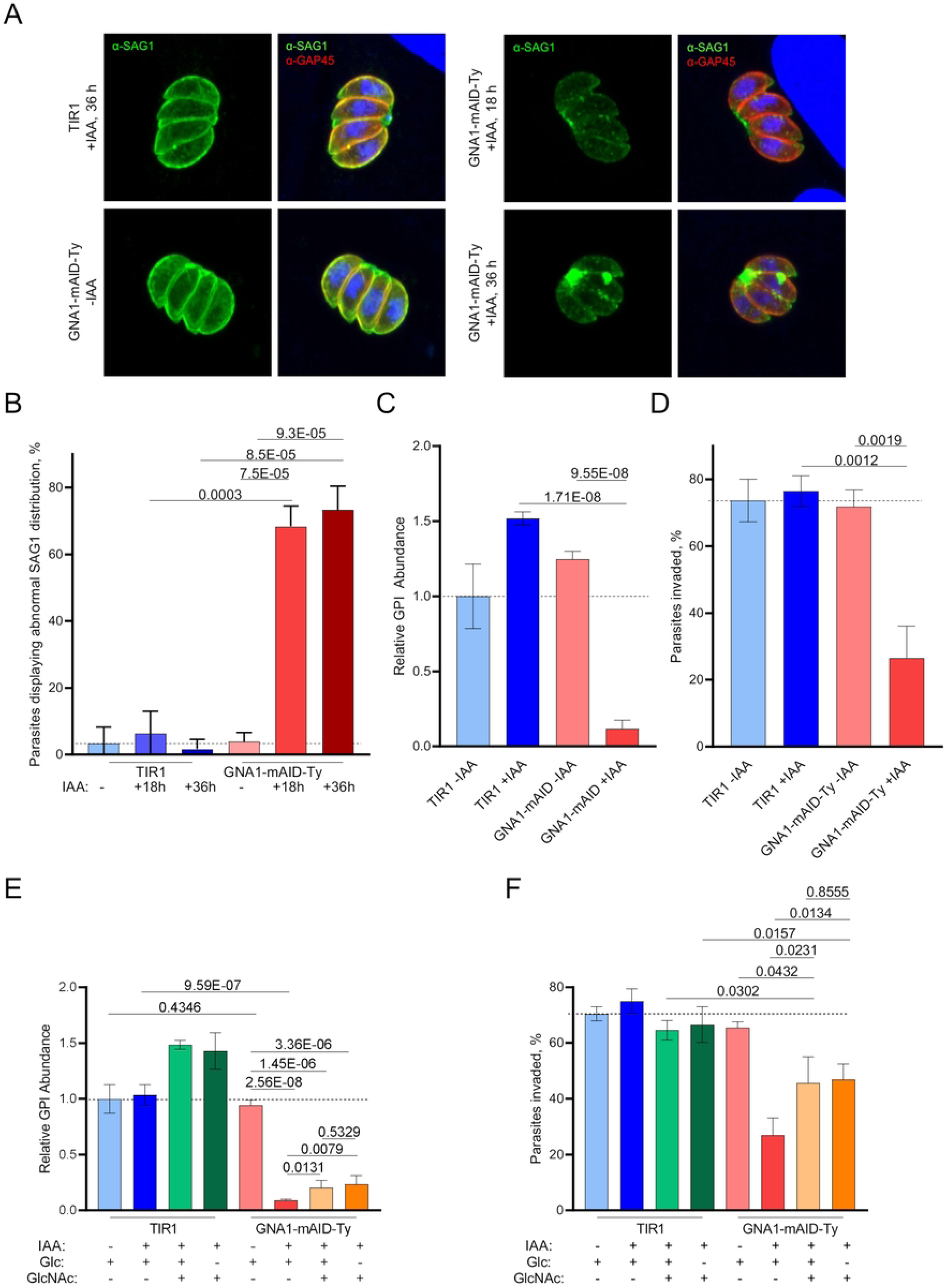
Lack of *Tg*GNA1 disrupts the localization of GPI-anchored proteins causing an invasion defect. **B)** Immunofluorescence assay (IFA), showing the staining of the pellicle marker GAP45 and the GPI-anchored protein surface antigen 1 (SAG1) in TIR1 and GNA-mAID-Ty parasites after varying durations of auxin (IAA) treatment. **B)** Quantification of vacuoles displaying normal (even distribution) and abnormal SAG1 signal (uneven, patchy distribution with predominant accumulation inside the residual body), based on IFA images as shown in panel A. **C)** relative GPI abundance of untreated (−IAA) or IAA- treated (+IAA, 36 h) TIR1 and GNA1-mAID-Ty parasite. **D)** Percent of invaded TIR1 and GNA1-mAID-Ty parasite in the absence (-IAA) or after IAA treatment (18 h) as determined by a red/green invasion assay. **E)** Quantification of GPI-anchors in cells grown with the indicated treatments/supplementations for 18 hours. **F)** Percentage of invaded TIR1 and GNA1-mAID-Ty parasites, following the indicated treatment over 18 hours. A) shows representative images of 3 independent experiments. B-F show representative data from one of three independent biological replicates, averaging technical triplicates. For B, >100 vacuoles were counted and categorised, per replicate. p-values are given in B-F following two-sided student’s t-tests, comparing the indicated conditions. Abbreviations: see Fig 3 and 4.

To assess if GlcNAc supplementation, and the consequent increase in UDP-GlcNAc levels under Glc-deplete conditions (Fig 4D), could restore GPI-anchor synthesis, GPI-anchor levels were quantified after GlcNAc supplementation under Glc-replete or - deplete conditions. After 18 hours of IAA-treatment and the indicated supplementations, GlcNAc supplementation appeared to increase relative GPI levels in TIR1 parasites and led to a slight increase in GPI levels in GNA1-depleted parasites. However, levels remained markedly lower (∼5-fold) compared to control conditions (Fig 5E). Notably, the modest increase in GPI-anchor levels was observed equally under Glc replete and Glc deplete conditions. These relative GPI levels correlated remarkably well with parasite invasion following 18 hours of treatment with IAA and supplementations as indicated: GlcNAc-supplemented GNA1-depleted parasites demonstrated a significant but still incomplete rescue in their ability to invade host cells (Fig 5F).

Overall, GlcNAc supplementation under Glc deplete conditions fully restored UDP-GlcNAc levels in cells lacking GNA1 (Fig 4D), but this only facilitated a modest and incomplete rescue of the intracellular growth rate (Fig 4E), GPI abundance (Fig 5E) and the parasites’ ability to invade (Fig 5F). In summary, GNA1 is essential for *T. gondii* replication and invasion and cannot be bypassed by GlcNAc salvage.

## Discussion

The amino sugar pathway, also known as the hexosamine biosynthetic pathway, plays a crucial role in various organisms, including the apicomplexan parasite *P. falciparum* [20, 21]. A critical function of the pathway in the related apicomplexan *T. gondii* is consistent with the highly negative fitness scores for most of the enzymes in the pathway, as reported by a genome-wide CRISPR sgRNA-based fitness screen [23]. However, the screen assigned a positive fitness score to GNA1, which catalyses the acetylation of GlcN6P to GlcNAc6P, contrasting with the assumed key function of this enzyme in the pathway. Our presented data confirm that the specific activity of GNA1 in *T. gondii*, as previously illustrated by *in vitro* activity assays [20], is essential for parasite survival. The discrepancy between our findings and the genome-wide fitness analysis likely arises from an omission in the gene annotation, which failed to highlight the existence of several short GNA1 isoforms. Our data suggest that full size GNA1 (consistent with the annotated sequence [25]) is synthesised alongside several shorter isoforms. These shorter isoforms contain the essential acetyltransferase domain [39], and remain unaffected by several individual sgRNAs employed in the genome-wide fitness screen to disrupt GNA1 [23]. To our knowledge, most eukaryotic organisms exhibit only a single GNA1 isoform [40]. However, within the apicomplexan GNA1 family, *T. gondii* GNA1 stands out due to its distinctive and elongated N-terminus [20]. The function of this extended N-term remains unknown. Despite the valuable information provided by genome wide studies, our findings highlight the limitations of such approaches and automatic gene annotation, reinforcing the importance of studying genes individually for a thorough comprehension of their significance.

UDP-GlcNAc and UDP-GalNAc derived from UDP-GlcNAc through the activity of GalE epimerase [7], are the main products of the amino sugar pathway. UDP-GlcNAc is key for the synthesis of GPI anchors and free GIPLs, which are present on the surface of all *T. gondii* life stages [41-43]. SAG1, the primary surface antigen of *T. gondii*, is a GPI anchored protein crucial for host cell binding and invasion [44]. Consequently GPI anchors and GIPLs contribute to parasite virulence [45] and are essential for *T. gondii* survival [16]. Our results reveal that GNA1 depletion leads to a marked drop in GPI anchors, altering the localization of SAG1 and severely impacting host cell invasion and intracellular growth. Furthermore, UDP-GlcNAc, along with other glycosylations, plays a crucial role in the biosynthesis of *N*-glycans, modifying numerous proteins in the *T. gondii* secretory pathway [7]. Several studies suggest that *N*-glycosylation is essential for parasite invasion, motility, and viability [8, 23, 46]. Indeed, *N*-glycosylation, but also GPI anchor biosynthesis and the amino sugar metabolism are among the metabolic pathways with the highest proportion of essential genes in *T. gondii* tachyzoites, as reported by a recent study [22]. In summary, our data emphasize the importance of GNA1 for the amino sugar pathway and UDP-GlcNAc synthesis, highlighting the pivotal role of this metabolic route for parasite virulence and survival.

Depletion of GNA1 results in the accumulation of GlcN6P, and the reduction or absence of the downstream metabolites GlcNAc6P, GlcNAc1P and UDP-GlcNAc. While growth can be rescued by supplementing the media with high concentrations of GlcNAc in *P. falciparum* and other organisms [21, 32, 33], GlcNAc supplementation fails to rescue the lack of GNA1 in *T. gondii.* GlcN supplementation also proved ineffective in recovering parasite growth. Despite the inability to rescue parasite growth, the absence of Glc in the media enhances GlcNAc salvaging, replenishing UDP-GlcNAc levels. This strongly suggests a competition between Glc and GlcNAc at the uptake or phosphorylation processes. Nevertheless, the recovery of GPI anchors and parasite growth through GlcNAc salvage is only partial when Glc is absent. The low levels of Fru6P indicate incomplete gluconeogenesis via glutamine, the predominant carbon source for *T. gondii* in absence of Glc [47]. In addition, these scant amounts of Fru6P in GlcNAc-supplemented TIR1 parasites under Glc depletion, strongly suggest the lack of an amino sugar catabolic pathway in *T. gondii*. Notably, the pronounced accumulation of GlcN6P observed in *T. gondii* GNA1 mutants could also contribute to the limited rescue observed with GlcNAc supplementation [49, 50]. Intriguingly, the accumulation of GlcN6P correlated more closely with a drop in GPI levels and concomitant impairment of parasite invasion than UDP-GlcNAc levels, indicating a potentially toxic impact of this surge in GlcN6P. Indeed, glucosamine has been shown to interfere with *P. falciparum* asexual intraerythrocytic growth at high doses [47-49], and a comparable effect could be occurring in *T. gondii*. Regardless, the high concentration of GlcNAc needed for partial *T. gondii* growth recovery remains far from physiological levels [50], suggesting that the likelihood of rescuing GNA1-depleted parasites under physiological conditions is very remote. In summary, the incomplete rescue in GlcNAc supplemented media strongly suggests the inability of a metabolic bypass to overcome GNA1 deficiency. This spotlights *T. gondii* GNA1 as a potential drug target to tackle toxoplasmosis.

*T. gondii* GNA1 belongs to a specific gene family, with an independent evolutionary origin within the phylum Apicomplexa [20]. Apicomplexan GNA1s exhibit distinct features and conserved motifs, and a recent structural study highlighted the divergent binding sites for GlcN6P and acetyl-CoA in *Cryptosporidium parvum* GNA1 compared to human GNA1, including important variations in key residues [21]. The key role of the amino sugar pathway for *T. gondii* viability, and the predicted significance of GPI anchors and GlcNAc-containing glycoconjugates across *T. gondii*’s life cycle [7, 22], underscore the potential of GNA1 as a versatile multistage therapeutic target in toxoplasmosis that could be exploited for selective parasite inhibition.

Current drug therapies for human toxoplasmosis lack specificity, often leading to adverse effects and inconsistent efficacy [5, 51]. Novel treatments against *T. gondii* must target the slow growing bradyzoites [52] to eradicate the chronic stage, which poses a threat to infected individuals if the immune system is compromised [53]. Targeting bradyzoites efficiently is hindered by several hurdles: drugs must cross the blood-brain barrier and traverse the cyst wall and must act on enzymes/pathways that are critical for the poorly characterized metabolism of bradyzoites [54]. Intriguingly, the cyst in which bradyzoites reside and persist is heavily glycosylated, containing high levels of GlcNAc and GalNac residues [55]. A previous study highlighted that glycosylation of the cyst wall is critical for *T. gondii* persistence [17]. Specifically, Caffaro et al., demonstrated that the nucleotide sugar transporter *Tg*NST1 is required for cyst wall glycosylation and its disruption impairs the ability of *T. gondii* to persist but is dispensable for tachyzoites *in vitro* and during acute infection *in vivo* [17]. We demonstrate here that GNA1 is essential for tachyzoites and can be expected to be essential for bradyzoites given the high need for UDP-GlcNAc and UDP-GalNAc during persistence [17], making it a promising candidate for a drug target.

## Material and Methods

### Parasite lines, culture and treatments

Parasites stably expressing TIR1 were a generous gift from the laboratory of David Sibley [29]. These were maintained by regular passages in human foreskin fibroblasts (HFF-1, ATCC SCRC-1041), in Dulbecco Modified Eagle Medium (DMEM, Gibco, 41966-029) supplemented with foetal bovine serum (FBS, Gibco, 10270-106, 5% v/v), L-glutamine (Gibco, 20530-024, additional 2 mM) and Gentamycin (Gibco, 15750-045, 25 μg/ml), incubated in humidified incubators at 37 °C and 5% CO2.

Auxin (IAA, Sigma-Aldrich, I-2886) was added to cultures at 500 μM final in ethanol as indicated for each experiment. Supplementations with sugars (Glc – Agilent, 103577, GlcN – Sigma-Aldrich, G1514 or GlcNAc – Sigma-Aldrich, A3286) were performed as described for each experiment in regular medium, as above or in DMEM without Glc (Gibco, 11966-025) supplemented with 5% (v/v) dialysed FBS (Pan Biotech P30-2102; 10,000 Da exclusion size membrane) and additional 10 mM glutamine (Agilent, 103579).

### Generation of Transgenic Parasites

The GNA1-mAID-Ty parasite line was generated through co-transfection of a CRISPR-Cas9 expression plasmid [27] with a guide RNA (P1, S2 Table) targeting the 3’-UTR of GNA1 (TTGT1_243600) and a homology repair template encoding the mAID domain, the 3-Ty domain and the *hxgprt* resistance cassette, amplified with the primers P2 and P3 (S2 Table) by KOD PCR (Sigma-Aldrich). Transfected parasites were selected in medium containing mycophenolic acid (25 μg/ml) and xanthine (50 μg/ml) over one week and cloned by serial dilution followed by a second round of cloning. Subclones were frozen and a single clone was used in the following experiments.

Correct integration of the homology template at the desired location was assessed by PCR (GoTaq DNA Polymerase, Promega) on extracted genomic DNA (Promega Wizard DNA Extraction) testing 3 amplifications using primers P4 and P5, P4 and P6 and P4 and P7 amplifying under the following conditions: 95 °C, 2 min; (95 °C, 15 s; 57 °C 15 s; 72 °C 1.5 min) × 35; 72 °C, 5 min on a SimpliAmp Thermal Cycler (Applied Biosystems).

### Immunofluorescence Assays

Confluent monolayer of HFF cells grown on coverslips were inoculated with 10 μl of freshly egressed parasite cultures and treated with IAA as or other supplementations as indicated for each experiment. Twenty-four hours after inoculation, parasites were fixed with 4% PFA and 0.05% glutaraldehyde for 10 min, before quenching with 0.1 M glycine in PBS for 20 min. Infected host cells were permeabilized using 0.2% Triton X-100/PBS for 20 min, followed by 20 min incubation in (2% BSA/0.2% Triton X-100/PBS to block unspecific binding and subsequently probed with different primary antibodies diluted in 2% BSA/0.2% Triton X-100/PBS for 1 hour. The following primary antibodies were used as indicated for each experiment: α-Ty (1:10, mouse monoclonal, BB2), α-SAG1 (1:10, mouse, T4-1E5), polyclonal rabbit α-GAP45 (1:10,000, used for growth assay) [56], monoclonal mouse α-actin (1:20) [57], polyclonal rabbit α-CPN60 [58] and mouse monoclonal α-5F4 (F1 ATPase beta subunit, P. Bradley). The probed monolayer was washed (3 × 5 min, 0.2% Triton X-100/PBS) and probed with a secondary antibody: anti mouse Alexa fluor 488 (Invitrogen, A11001), anti-rabbit Alexa fluor 594 (Invitrogen, A11012). Following 3 washing steps as above, the coverslips were mounted on microscopy slides using DAPI-containing FluoromountG (SouthernBiotech). Slides were viewed on an Eclipse Ti inverted microscope (Nikon). For growth assays, the number of parasites was counted in >100 vacuoles per condition for 3 independent biological replicates. Images were acquired using an LSM 700 confocal scanning microscope (Zeiss) and images were processed using Fiji Image J software.

### Western Blots

Parasites were harvested from a freshly lysed dish, washed with PBS, and resuspended in SDS–PAGE buffer (50 mM Tris-HCl, pH 6.8, 10% glycerol, 2 mM EDTA, 2% SDS, 0.05% bromophenol blue, and 100 mM dithiothreitol (DTT)). Following boiling for 10 min, samples were subjected to SDS–PAGE under reducing conditions. Proteins were transferred to a hybond ECL nitrocellulose membrane using a wet transfer system (Bio-Rad Laboratories, Hercules, CA, USA). The membrane was incubated in α-Ty antibody (1:10, mouse monoclonal, BB2) and rabbit α-catalase as a loading control [59], diluted in PBS, 0.05% Tween20, 5% skimmed milk. Following 3 washing steps, the membrane was incubated with the secondary antibodies (goat α-mouse, horse radish peroxidase conjugated, Sigma Aldrich, A5278). Signal was visualized using the SuperSignal West Pico PLUS Chemiluminescent Substrate (ThermoFisher Scientific, 34580). Images were taken using the Bio-Rad ChemiDoc MP Imaging System and images were processed using Bio-Rad Image Lab software.

### Plaque Assays

Serial dilutions of parasite cultures were incubated on a confluent host cells monolayer in 12- or 24-well plates for 7 days. Afterwards, the infected monolayer was washed with PBS and fixed with 4% paraformaldehyde (PFA) for 10 min. Host cells were stained with a crystal violet solution (12.5 g crystal violet, 125 ml ethanol mixed with 500 ml water containing 1% (*w*/*v*) ammonium oxalate) over 3 hours. Wells were washed 3 times with deionized water to remove excess crystal violet and images of dried wells recorded.

### Invasion (Red/Green) Assays

Parasites from a freshly egressed culture treated as described for each experiment were diluted 1:10 and 150 μl of parasite solution used to infect a coverslip with confluent HFFs in a 24-well plate. The plate was gently spun for 1 min at 1,100 *g* and subsequently incubated in a water bath at 37 °C. Cells were fixed with 4% PFA and 0.05% glutaraldehyde for 7 min, before quenching with 0.1 M glycine in PBS for 10 min. Unspecific binding was blocked with (2% BSA in PBS – without triton), followed by incubation with α-SAG1 (1:10, mouse, T4-1E5) as above but without triton. Wells were washed 3 times with PBS before fixing cells with 4% PFA for 7 min. The next steps, permeabilization, blocking, primary antibody incubation, washing, secondary antibody incubation, washing and mounting were carried out as described above for the IFA. Polyclonal rabbit α-GAP45 (1:10,000) [56] was used as primary antibody and SAG1 and GAP45 were revealed in green and red, respectively, using the secondary antibodies as above for the IFA. Slides were viewed on an Eclipse Ti inverted microscope (Nikon). More than 100 parasites were counted per biological triplicate and categorised as invaded (red staining only) or non-invaded (red and green staining).

### Harvest of parasites for mass spectrometry analyses

Freshly egressing or intracellular parasites were harvested as follows: medium was aspirated, and the metabolism quenched through addition of ice-cold PBS. The monolayer was scraped, and parasites released via multiple passages through a 26G needle. The parasite solution was passed through a filter of 3 μm exclusion size (Merck-Millipore, TSTP04700) to remove host cell debris and collected in 15 ml conical tubes. Parasites were pelleted (2000 *g*, 4 °C, 25 min) and washed two more times with ice-cold PBS. Residual PBS was removed, and pellets of 10^8^ parasites resuspended in medium as indicated below for labelling of extracellular parasites or stored at ™80 °C until metabolite extraction.

### Stable Isotope Labelling of Extracellular Parasites

Parasites pellets were resuspended in 2 ml of DMEM without Glc (Gibco, 11966-025) supplemented with 5% (v/v) dialysed FBS (Pan Biotech P30-2102; 10,000 Da exclusion size membrane) and 10 mM U-^13^C6-Glc (Cambridge Isotope Laboratories, CLM-1396) or U-^13^C6-glucosamine (Cambridge Isotope Laboratories, CLM-9883) or U-^13^C6-*N*- acetylglucosamine (Cambridge Isotope Laboratories, CLM-1827) and incubated in a conical tube for 5 hours in humidified incubators at 37 °C and 5% CO2 prior to addition of excess ice-cold PBS, centrifugation and PBS washes as described above.

### Sample Preparation for GC-MS Analyses

Metabolite extraction and derivatization was performed as previously described but without heating step [60]. In brief, parasite pellets were placed on ice for 5 min before addition of 50 μl chloroform followed by 200 μl methanol:ultrapure water (3:1, including scyllo inositol as an internal standard, 1 nmol, Sigma-Aldrich, I8132). Extraction was facilitated through vigorous vortexing. Samples were spun (20,000 *g*, 4 °C, 10 min) and the supernatant transferred to a new vial containing 100 μl ice-cold ultrapure water. Samples were vortexed and spun (20,000 *g*, 4 °C, 10 min). The lower, organic phase (apolar, 50 μl) and the upper, polar phase (300 μl) were processed further as outlined below.

### Sample Preparation for LC-MS Analyses

Cells were harvested and washed as described above. Pellets were reconstituted in 60 μl acetonitrile:ultrapure water (4:1, containing ^13^C6/^15^N-isoleucine as internal standard, 40 μM, Cambridge Isotope Laboratories, CNLM-561-H) and vortexed vigorously. Extracts were spun (20,000 *g*, 4 °C, 10 min) and the clear supernatant transferred to a mass spectrometry vial with insert. The metabolite extraction is based on that described in previous studies [61].

### GPI Quantification via GC-MS

GPI quantification analysis and quantification was performed via methanolysis as previously described [38, 62]. The apolar phase was transferred to a flame-sealed glass tube (Sigma-Aldrich, Z328510) and dried in a centrifugal evaporator. Next, 50 μl methanolic hydrochloric acid (HCl, Supelco, 33354) were added, the tube flame-sealed under vacuum and incubated in an oven at 80 °C over night. The next day, the glass tube was opened, and the content transferred to a mass spectrometry vial insert containing 10 μl pyridine to neutralise the pH. The solution was dried in a centrifugal evaporator and further derivatised through addition of 20 μl pyridine and 20 μl N, O- Bis(trimethylsilyl) trifluoracetamid 99% (Supelco, B-023). Samples were analysed on an 8890 GC System (Agilent) equipped with a DB5 capillary column (J&W Scientific, 30 m, 250 μm inner diameter, 0.25-μm film thickness), with a 10-m inert duraguard, connected to a 5977B GC/MSD in electron impact (EI) mode equipped with 7693A autosampler (Agilent). The GC-MS settings were as follows: Inlet temperature: 270 °C, MS transfer line temperature: 280 °C, MS source temperature: 230 °C and MS quadrupole temperature: 150 °C. The oven gradient during the sample run was as follows: 80 °C (2 min); 80 °C to 140 °C at 30 °C/min; 140 °C to 250 °C at 5 °C/min; 250 °C to 310 °C at 15 °C/min; 310 °C for 2 min. Mannose and palmitic acid derivatives were identified based on the ion spectrum and retention time of authentic standards. Final analyses were performed in selected ion monitoring (SIM) mode, detecting the ions *m/z* 204 (mannose derivative) and *m/z* 270 (palmitate derivative), following injection of 1 μl in split mode (1:50). Data was analysed using MassHunter (Quantitative Analysis and Qualitative Analysis 10.0, Agilent) and Excel (Microsoft).

### Untargeted Polar Metabolite Profiling via GC-MS

Polar metabolites were derivatised and analysed as previously described [60]. In brief, the polar phase was sequentially dried within a mass spectrometry insert in a centrifugal evaporator (50 μl at a time) and further dried and concentrated through addition of methanol. The dried metabolite extract was derivatised through addition of 20 μl pyridine containing methoxyamine hydrochloride at 20 mg/ml and incubation at room temperature overnight. The following day, 20 μl N, O-Bis(trimethylsilyl) trifluoracetamid 99% (Supelco, B-023) were added and samples vortexed and analysed by GC-MS. The analysis was performed as described above but using the following oven gradient: 70 °C (1 min); 70°C to 295 °C at 12.5 °C/min; 295 °C to 320 °C at 25 °C/min; 320 °C for 2 min and operating in scan mode (*m/z* 70-700) with a 5.5 min solvent delay. Metabolites were identified based on the analysis of authentic standards or reliable predictions (NIST library, NIST MS Search 2.4, >60% confidence and manual curation). Data was analysed using MassHunter (Quantitative Analysis and Qualitative Analysis 10.0, Agilent) and Excel (Microsoft). Metabolite intensities were normalised to the internal standard (scyllo inositol) and expressed as relative abundances relative to the control (TIR1 −IAA, abundance = 1).

### Targeted Aminosugar Profiling via GC-MS

Samples were prepped and analysed as described above for the untargeted profiling. However, the MS was operated in SIM mode, detecting the ions *m/z* 356, 357, 358 and 359 to determine labelling in the desired sugars (see S3 Fig), as well as *m/z* 318 (internal standard, scyllo inositol). The ^13^C-fractional labelling was determined by measuring the isotopologue abundance for *m/z* 357, 358 and 359 and correcting for occurrence of natural isotopes [63].

### Targeted Aminosugar Profiling via LC-MS/MS

Sample analyses were performed on an Agilent LC-MS (Santa Clara, CA, USA) using MassHunter B.08.00 software for system control and data acquisition. 1290 Infinity LC comprised a binary pump, HiP autosampler, column oven, and Flexible Cube module. The LC was hyphenated to a 6490 triple-quadrupole detector through an Agilent Jet Stream ion source. HILIC chromatographic separation was conducted on a Waters Acquity Premier BEH Amide column (2.1 × 150 mm, 1.7 µm) kept at 35 °C. Elution was performed at a flow rate of 0.4 ml min^−1^, using the following gradient of mobile phases [64]: A (10 mM AF + 0.15% FA in MeCN:H2O 85:15 v/v) and B (10 mM AF + 0.15% FA in H2O): 0-6 min 0% B, 6.1 min 5.9% B, 10 min 17.6% B, 12 min 29.4% B and back to 0% B from 12 to 18 minutes for column re-equilibration. Samples were kept at 6 °C and injection volume was 7 µl. Ion source parameters were as follow: Jet Stream gas temperature and flow rate were 250 °C and 15 l min^−1^ respectively, while for sheath gas they were set to 400 °C and 11 l min^−1^. Nebulizer pressure was 40 psi, and a 3000 V capillary voltage was used. Ion funnel high/low pressure radiofrequencies were set to 150/60 for positive ionization transitions and 90/60 for negative ones.

Multiple reaction monitoring transitions were optimized using Agilent MassHunter Optimizer B.08.00 using individual solutions of the compounds dissolved at 100 μM in 80% ACN. Collision energy and cone voltage were optimized and at least three fragments derived from the [M+H]^+^, [M+Na]^+^ or [M-H]^−^ precursor ions were used to monitor each molecule. The specific transitions, MS source and ion funnel conditions can be found in S8 Table. Data was processed using Skyline 22.2. Peak identity was confirmed based on its qualifier transitions and retention time compared to those of standard compounds (RSD < 3%). The relative abundance of each molecule was expressed as the sum of all the areas of the corresponding transitions, normalized to the internal standard and expressed as relative abundance in relation to TIR1 −IAA abundance = 1, using Excel (Microsoft).

## Author contributions

J.K., L.I., D.S.F. and M.P.A. conceived the study; M.P.A. and J.K. designed, performed and interpreted the experimental work, with the support of L.I. and D.S.F; J.K. conducted the formal analysis; V.G.R. and S.R. were responsible of LC-MS experiments; J.K., L.I. and D.S.F. supervised the research; D.S.F contributed to resources; M.P.A. and J.K. outlined the draft. All authors contributed to the writing, review and editing of this manuscript.

## Author Summary

*Toxoplasma gondii*, *Plasmodium*, and *Cryptosporidium* spp., pose serious threats to human health. *T. gondii*, an intracellular and opportunistic pathogen, cunningly avoids the host immune defences by forming long-lasting tissue cysts. Finding effective drugs to eliminate these parasites remains a challenge.

The glucosamine-phosphate-*N*-acetyltransferase (GNA1) catalyses a critical key step in the production of activated sugar nucleotides to build glycoconjugates essential for various functions in the cell. In *P. falciparum*, this enzyme has been identified as a potential target for antimalarial drugs.

In this study, we explored the importance of this pathway in *T. gondii* and discovered that these sugar-containing compounds play a vital role in the parasite’s ability to invade and replicate in host cells – crucial processes for its survival and ability to cause disease. Intriguingly, unlike some organisms that can bypass the pathway, *T. gondii* relies critically on glucosamine-6-phosphate acetylation. This reliance sheds light on the parasite’s distinct metabolic properties and highlights the pathway’s potential as a target for new therapeutic strategies.

## Captions of Supporting Material

**S1 Fig. GNA1 sequence, genome-wide fitness screen guides, locus modification and genomic PCR. A)** GNA1 coding sequence as found on ToxoDB. The initial start codon is highlighted in green, the stop codon in red. Four additional ‘in-frame’ start codons were found and are also highlighted by green shading. Blue (guide with positive score) and red shading (guide with negative score; intensity of shading indicating score) highlights the sequence of single guide RNAs (sgRNA) used in the genome-wide fitness screen. For overlapping guides, the first guide is shown as underlined, the second guide is shown in bold. The name/number of the guides and their respective fitness score is provided. The listing is from left to right and from top to bottom in the order of their appearance in the coding sequence. Guides with a positive fitness score are shown in blue, guides with a negative fitness score in red. Note that the first 4 guides only affect the longest putative GNA1 product, while the last 4 guides affect all GNA1 products, including the shortest potential GNA1 protein (highlighted in italic). The GNAT domain, needed for the catalytic activity is highlighted in purple and bold. **B)** Schematic depiction of the GNA1 locus and its modification through insertion of a mini auxin inducible degron (mAID) domain, a 3-Ty tag and a hxgprt cassette for selection. **C)** Schematic showing the binding sites of primers used to validate the successful modification of the GNA1 locus and integration PCR, showing the expected bands following amplification with the indicated primers. Sequences of these primers are listed in Supplementary Table 1.

**S2 Table. Description and sequence of primers used in this study.**

**S3 Fig. Chromatogram, ion spectrum and structure of relevant metabolites. A)** Authentic standards of glucose-6-phosphate (Glc6P), glucosamine-6-phosphate (GlcN6P) and *N*-acetylglucosamine-6-phosphate (GlcNAc6P) were derivatised via methoximation and silylated. Their gas chromatography-mass spectrometry (GC-MS) chromatograms were overlayed (top panel) and the corresponding ion spectra at the indicated retention times (RT) are shown below. Note that all 3 derivatives share a common fragment of *m/z* 357 (highlighted in red), which contains 2 carbons of the sugars. **B)** The structure of the methoxyamine (MeOx) and trimethylsilyl (TMS) derivative of ^13^C6-GlcNAc6P is shown. Labelled carbons are highlighted through red asterisks. The fragment *m/z* 357 (natural abundance) becomes *m/z* 359 in ^13^C-labelled sugars.

**S4 Fig. Untargeted metabolite profiling by gas chromatography-mass spectrometry (GC-MS).** Relative levels of 64 metabolites were determined in TIR1 and GNA1-mAID-Ty parasites in presence (+IAA, 36 hours) or absence of auxin (−IAA). Metabolites were quantified in equal cell numbers, normalised to an internal standard, and are shown relative to levels in TIR1 −IAA (level of metabolites TIR1 −IAA = 1). Metabolites were grouped into different categories. The fold-change is indicated through red and blue shading as shown in the legend. Significantly altered levels are highlighted through bold lettering. Data represent average of 4 independent biological replicates from a single experiment. Significance indicates a p-value <0.05 in a two-sided student t-tests in comparison to TIR1 −IAA.

**S5 Fig. Detail of amino sugar abundance graph** Detail of graph shown in main Fig 4D.

**S6 Fig Plaque assay following supplementation with varying glucose/GlcNAc ratios.** Plaque assay of TIR1 and GNA1-mAID-Ty parasites in presence or absence of auxin (IAA) and with glucose (Glc) or *N*-acetylglucosamine (GlcNAC) supplemented in medium without Glc as indicated. Image of stained plaques of a single experiment.

**S7 Fig. Morphology of the apicoplast and mitochondrion following downregulation of GNA1.** Immunofluorescence assays (IFAs) were performed after auxin (IAA) treatment for the indicated duration and after 24 hours of intracellular growth. Cells were stained with antibodies marking actin (Act) and the pellicle (GAP45), while organelles were stained with CPN60 (apicoplast) and 5F4 (mitochondrion). Intactness of the apicoplast and mitochondrion was determined in 3 technical replicates of a single experiment. Images show representatively the morphology of the apicoplast and mitochondrion.

**S8 Table. Transitions, MS source and ion funnel conditions.** Table with details pertaining to LC-MS analyses.

